# Katanin is involved in Microtubule Polymerization into Dendritic Spines and regulates Synaptic Plasticity

**DOI:** 10.1101/2022.05.04.490623

**Authors:** Franco L. Lombino, Jürgen R. Schwarz, Yvonne Pechmann, Michaela Schweizer, Markus Glatzel, Christine E. Gee, Kira V. Gromova, Matthias Kneussel

## Abstract

Dynamic microtubules transiently polymerize into dendritic spines, however intracellular factors that regulate this process and their functional role at synapses are hardly understood. Using live imaging, electrophysiology, and glutamate uncaging, we show that the microtubule-severing complex katanin is located at individual spine synapses, participates in the activity-dependent process of microtubule polymerization into dendritic spines, and regulates synaptic plasticity. Overexpression of a dominant-negative ATPase-deficient katanin subunit, did not alter microtubule growth velocities or comet density in dendrites, but significantly reduced the activity-dependent invasion of microtubules into dendritic spines. Notably, functional inhibition of katanin significantly affected the potentiation of AMPA-receptor-mediated excitatory currents after chemical induction of long-term potentiation (cLTP). Furthermore, interference with katanin function prevented structural spine remodeling following single spine glutamate uncaging. Our data identify katanin at individual spine synapses in association with PSD-95. Thus, katanin regulates postsynaptic microtubules and modulates synaptic structure and function.

## Introduction

Microtubules (MTs) are polymers comprised of α- and β-tubulin dimers that assemble in a head-to-tail fashion, resulting in filaments with plus and minus ends. At their plus-ends MTs are highly dynamic, polymerizing and depolymerizing through a stochastic process known as dynamic instability (1). While in most cells MTs are anchored near the centrosome, in neurons MTs can be nucleated in neurite projections, for instance at branch points following local MT severing (2). Polymeric MTs are transported by motor proteins into axons or dendrites (3) and MTs themselves serve as tracks for cargo transport by motor proteins (4–9). Whereas kinesin family proteins (KIFs) transport cargo towards MT plus-ends, dynein mediates minus-end-directed transport. Notably, MT plus ends are oriented distally in axons, while anti-parallel orientations are reported in dendrites (10). Neuronal MT transport is sensitive to tubulin posttranslational modifications (11, 12) and to microtubule-associated proteins (MAPs) (13, 14), both of which can regulate MT binding affinities and/or MT stability, respectively.

In contrast to the high abundance of MTs in dendrites (15), dendritic spines are typically rich in actin filaments (16–18). Visualization of the microtubule +TIP protein EB3 revealed however, that MTs in distal dendrites are very dynamic and occasionally polymerize into dendritic spines (19, 20). Since MT-associated transport is 5-10 times faster than MT polymerization, even transient MT invasion of spines would likely deliver cargo to post-synapses (9). Indeed, MTs growing into dendritic spines have been shown to transport KIF1A and its cargo synaptotagmin-IV to postsynaptic sites (21). MT entries into spine protrusions last seconds to minutes, following NMDA-type glutamate receptor (NMDAR) activation and calcium influx (22–24). Notably, chemically-induced long-term potentiation (LTP) increases MT entries into spines, whereas chemically-induced long-term depression (LTD) decreases them (22–24).

Together with polymerization and depolymerization MT severing contributes to sculpting and dynamically reorganizing MT arrays (2, 25). The three major MT severing proteins are the AAA-ATPases spastin, katanin and fidgetin (2, 26). Katanin functions as a hexameric assembly of dimers made of 60 kDa catalytic ATPase subunits with 80 kDa regulatory subunits (2, 27). In addition to an important role in cell division and adult neurogenesis (28), mammalian katanin regulates the number and length of neuronal MTs (26). It further interacts with CAMSAPs to regulate the organization and stability of MT minus ends (29). Notably, the MT-associated protein tau protects MTs from katanin-mediated severing in axons (30).

Here, we aimed to investigate mechanistic insights into microtubule severing at synapses. To this end, we asked whether katanin is located at dendritic spines and/or synaptic sites and may interact with the postsynaptic density (PSD). We further asked whether dominant-negative interference with katanin-mediated severing, affects MT polymerization into spines or alters synaptic structure and/or function. Our data connect microtubule severing with the regulation of neuronal plasticity.

## Results

### Katanin is located at dendritic spine synapses and associates with PSD-95

The role of katanin is closely related to microtubule (MT) severing in the regulation of a dynamic MT cytoskeleton (2, 26). Since MTs frequently grow into dendritic spines in an activity-dependent manner (19, 20), we asked whether katanin might also be located at spine protrusions. Dendrites and F-actin-positive spines stained positive for the endogenous catalytic subunit katanin p60 (Figure S1A-E) and the endogenous regulatory subunit katanin p80 (Figure S1F-J), with about 25% of all spines containing p60 (Figure 2E, left). Katanin p60 colocalized with the postsynaptic AMPA receptor (AMPAR) subunit GluA2 in apposition to synaptophysin-positive presynaptic terminals (Figure 1A-D and FigureS2A). Likewise, the regulatory katanin subunit p80 was found in colocalization with GluA2 (Figure 1E-H and Figure S2B), indicating that the MT severing complex is found at individual excitatory postsynaptic sites.

**Figure 1.**
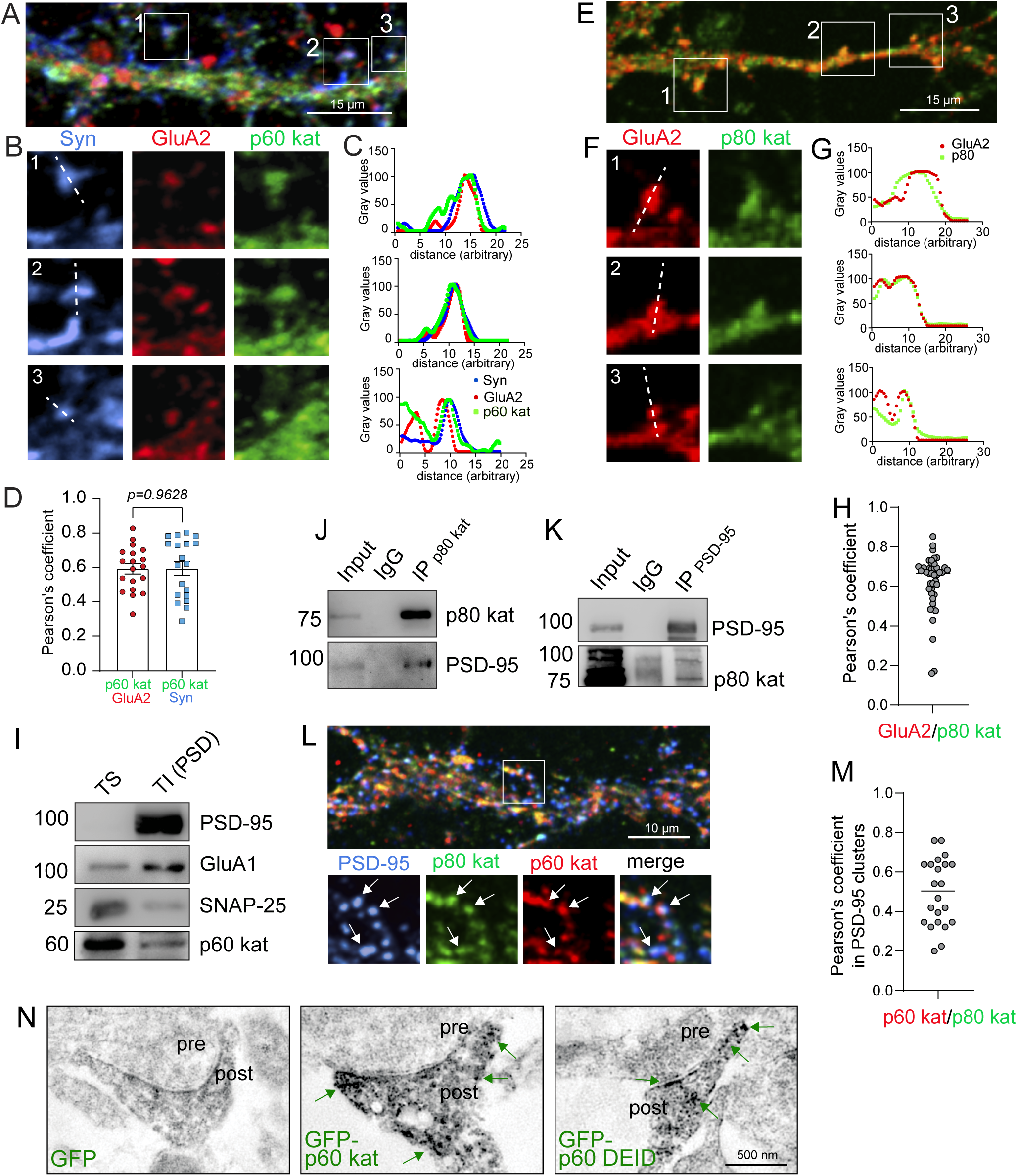
Katanin is located at glutamatergic spine synapses and binds to PSD-95. (A) Triple-immunostaining, endogenous katanin p60 (green), AMPAR subunit GluA2 (red), presynaptic marker synaptophysin (Syn, blue), DIV16-17 neurons, N = 3 experiments. (B) Magnification of boxed regions in A. (C) Profile plot of lines in B. (D) Pearson’s colocalization coefficient between p60 and GluA2 and p60 and Syn. p60/GluA2: Mean±S.E.M = 0.5918±0.02948, n = 19 ROIs; p60/Syn: Mean±S.E.M = 0.5942±0.03946, n = 19 ROIs. Mann-Whitney-U-test p = 0.8568. (E) Coimmunostaining, endogenous katanin p80 (green), AMPAR subunit GluA2 (red), DIV13-16 neurons, N = 3 experiments. (F) Magnification of boxed regions in E. (G) Profile plot of lines in F. (H) Pearson’s correlation coefficient of p80 katanin and GluA2. Mean±S.E.M = 0.6198±0.02192. n = 43 ROIs. (I) Western blot detection of katanin p60 in PSD fractions enriched for postsynaptic (PSD-95, GluA1) and containing very little presynaptic (SNAP-25) markers. TS: triton-soluble supernatant, TI (PSD): triton-insoluble postsynaptic density-enriched, (n = 3). (J) Co-IP with p80 katanin-specific antibody depicting precipitation of p80 katanin and co-precipitation of PSD-95 (n = 3). (K) Co-IP with PSD-95-specific antibodies depicting precipitation of PSD-95 and co-precipitation of p80 katanin (n = 3). (L) Triple immunostaining of p60 katanin (red), p80 katanin (green) and PSD-95 (blue) with magnification of triple colocalized puncta (M) Pearson’s correlation coefficient measured in PSD-95 enriched spots, Mean±S.E.M = 0.5218±0.01498, n = 22 ROIs. (N) DAB immunoelectron microscopy of dendritic spines of hippocampal cultures transfected either with GFP, EGFP-p60 katanin or EGFP-p60DEID katanin.

**Figure 2.**
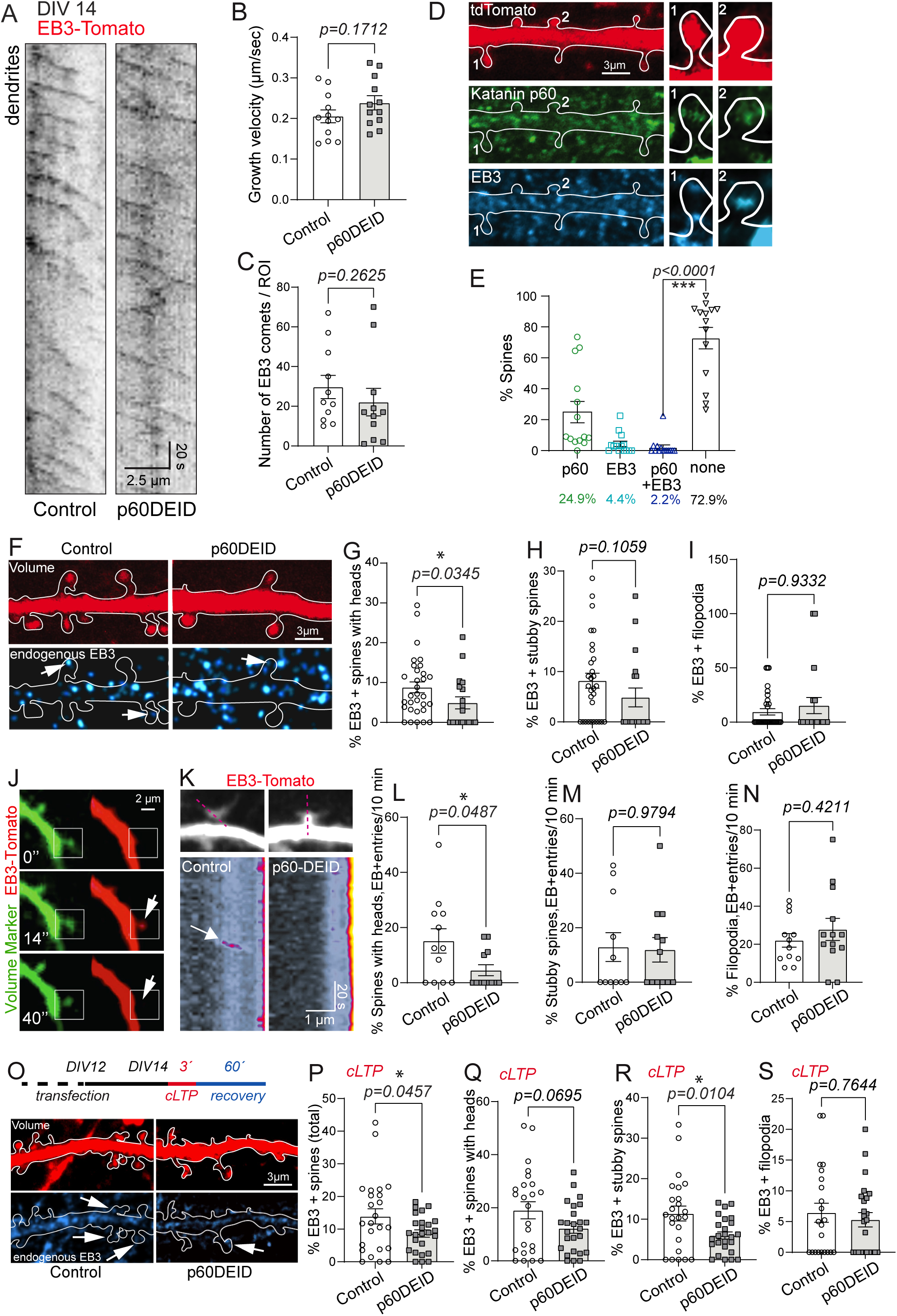
Functional inhibition of katanin reduces microtubule polymerization into dendritic spines. (A) EB3-tomato time-lapse imaging to visualize MT growth rates (EGFP control: left, EGFP-p60DEID, right). (B) Quantification of A shows comparable growth velocity among conditions. Control: 0.2054±0.01584 µm/s, p60DEID: 0.2387±0.01736 µm/s, Unpaired T-test, two tailed, p = 0.1712, (Control n = 12 ROIs, 503 comets); (p60DEID n = 12 ROIs, 340 comets). (C) Quantification of A shows comparable comet density among conditions. Control: 29.73±5.86, p60DEID: 22.09±6.93, Mann-Whitney-U-test p = 0.2625, (Control n = 12 ROIs, 503 comets); (p60DEID n = 12 ROIs, 340 comets). (D) Immunostaining of tdTomato expressing cells (volume marker), endogenous katanin p60 (green), endogenous EB3 (cyan), DIV12 neurons, N = 3 experiments. (E) Quantification of D. 24.94±6.94% spines contain p60 katanin. 4.42±1.76% spines contain EB3. 2.185±1.60% spines contain katanin and EB3. 72.82±6.96% spines contain neither katanin nor EB3, n = 14 ROIs; 440 spines. Comparison: % spines p60 katanin and EB3 positive vs. % spines p60 katanin and EB3 negative, Mann-Whitney-U-test, p<0.0001. (F-I) Immunostaining, endogenous EB3 (cyan), tdTomato (volume marker, red), DIV14 neurons, N = 3 experiments. (G) Quantification of % of EB3-positive spines with heads. Control: 8.81±1.35%, p60DEID: 4.90±1.53%, Mann-Whitney-U-test p = 0.0345 (Control: n = 33 cells, 979 spines; p60DEID: n = 20 cells, 402 spines). (H) Quantification of % of EB3-positive stubby spines. Control: 8.18±1.50%, p60DEID: 4.86±1.87%, Mann-Whitney-U-test, p = 0.1059 (Control: n = 33 cells, 505 spines; p60DEID: n = 20 cells, 187 spines). (I) Quantification of % of EB3-positive filopodia. Control: 9.52±2.90%, p60DEID: 15.26±7.43%, Mann-Whitney-U-test, p = 0.9332 (Control: n = 33 cells, 137 spines; p60DEID: n = 20 cells, 59 spines). (J) Time-lapse microscopy example of EB3 invasion into spines. Signal intensity was artificially increased to visualize the transient invasion and removal of the red EB3 signal (boxed region). (K-N) Time-lapse video microscopy using EB3-Tomato. (K) upper panels: EB3-tomato maximum projection depicts cellular morphology. Lower panels: Kymograph of EB3 +TIP invasion along depicted line in upper panels. N = 3 experiments. (L) Percentage of spines with heads invaded by EB3-Tomato in 10 minutes: Control: 15.16±4.430%, p60DEID: 4.54±2.01%, Mann-Whitney-U-test, p = 0.0487. (Control: n=12; p60DEID: n = 13 cells). (M) Percentage of stubby spines invaded by EB3-Tomato in 10 minutes: Control: 12.90±5.29%, p60DEID: 11.93±4.51%, Mann-Whitney-U-test, p = 0.9794. (Control: n = 12; p60DEID: n = 13 cells). (N) Percentage of filopodia invaded by EB3-Tomato in 10 minutes: Control: 22.07±3.58%, p60DEID: 27.78±5.82%, Unpaired two-tailed t-test p = 0.4211. (Control: n = 12; p60DEID: n = 13 cells). (O-S) Three minutes of chemical stimulation and recovery. Subsequent immunostaining, endogenous EB3 (cyan), tdTomato (volume marker, red), DIV12-14 neurons, N = 3 experiments. (P) Quantification of % of all EB3-positive spines after stimulation. Control: 13.91±2.34%, p60DEID: 8.65±1.11%, Unpaired two-tailed t-test p =0.0457 (Control n = 24 cells, 2558 spines; p60DEID n = 25 cells, 3174 spines). (Q) Quantification of % of EB3-positive spines with heads after stimulation. Control: 19.04±3.21%, p60DEID: 12.23±1.85%, Unpaired two-tailed t-test p = 0.0695 (Control: n = 24 cells, 1081 spines; p60DEID: n = 25 cells, 1472 spines). (R) Quantification of % of EB3-positive stubby spines after stimulation. Control: 11.39±1.85%, p60DEID: 5.98±0.88%, Unpaired two-tailed t-test p = 0.0104 (Control: n = 24 cells, 1040 spines; p60DEID: n = 25 cells, 1213 spines). (S) Quantification of % of EB3-positive filopodia after stimulation. Control: 6.44±1.55%, p60DEID: 5.30±1.17%, Mann-Whitney-U-test, p = 0.7644 (Control: n = 24 cells, 437 spines; p60DEID: n = 25 cells, 489 spines).

Differential centrifugation or a sucrose gradient to enrich postsynaptic densities (PSDs) confirmed this result. Katanin subunits were not only found in the supernatant, but were detected in plasma membrane-enriched P2 fractions (Figure S1K). Likewise, katanin p60 was detected at PSDs that were enriched for the postsynaptic proteins PSD-95 and GluA1, and contained little amounts of the presynaptic t-SNARE protein SNAP-25 (Figure 1I). We further performed co-immunoprecipitation from mouse forebrain lysate to investigate whether katanin associated with postsynaptic proteins. Strikingly, katanin p80- specific antibodies precipitated the p80 subunit and co-precipitated PSD-95 (Figure 1J). Vice versa, PSD-95-specific antibodies precipitated PSD-95 and co-precipitated p80 (Figure 1K). Triple immunostaining confirmed the colocalization of both katanin subunits with PSD-95 (Figure 1L-M and Figure S2C), indicating that the katanin complex associates with the postsynaptic scaffold at synapses. Furthermore, immunoelectron microscopy with diaminobenzidine (DAB) confirmed the postsynaptic localization of GFP-p60 katanin and a dominant-negative mutant GFP-p60DEID (27) at ultrastructural resolution in spines of cultured primary neurons (Figure 1N). These data demonstrate the presence of the microtubule severing complex katanin at spines in association with postsynaptic sites irrespective of its severing activity.

### Dominant-negative inhibition of katanin function reduces microtubule polymerization into dendritic spines

Two p60 katanin-like subunits are expressed in neurons and might associate with katanin hetero-oligomers (31, 32). Therefore, p60 depletion could be compensated by these alternative subunits. To interfere with katanin-mediated MT severing robustly, we overexpressed a validated dominant-negative katanin construct (katanin p60-DEID) known to associate with microtubules (27) (Figure S3K), which prevents ATP hydrolysis and consequently MT severing activity (27).

First, we asked whether functional katanin inhibition affects the growth rates of microtubules in neuronal dendrites by overexpressing p60DEID at neuronal stages DIV12-17 when dendritic spines are present (Figures 1, 2, Figures S1, S2). We observed no changes in either MT growth velocity or EB3 comet density (Figure 2A-C). In contrast, we detected significantly reduced microtubule growth velocities at DIV4, an early neuronal stage that is characterized by dendrite outgrowth (33), but still lacks spine protrusions (33) (Figure S3A-C). These data suggest that katanin might regulate microtubules during dendritogenesis but is not a major determinant of microtubule dynamics in mature neuronal dendrites containing spines.

Independent experiments analyzing endogenous EB3 in DIV 12-17 neurons by immunostaining revealed that a small percentage of dendritic spine protrusions (4.4% - 8.7%) contained EB3-positive microtubule +TIPs at a given time (19, 20) (Figure 2D, E, cyan and Figure S3D, left). About half of these spines were double-positive for endogenous EB3 and endogenous katanin p60 (Figure 2D, E). Therefore, we conclude that katanin can in general be present in spines containing dynamic MTs (compare with Figure 1N). Notably, functional inhibition of katanin-mediated microtubule severing (27) specifically reduced the percentage of those EB3-positive spines that contained a head (Figure 2F-I and Figure S3D), while the overall percentage of different spine types (Figure S3E-G) and the amount of EB3 in dendrites and spines (Figure S3H) remained unaltered. These data suggest that functional inhibition of katanin mainly affects MT growth towards mature spines, which are typically characterized by abundant amounts of AMPARs and high glutamate sensitivity (34).

Using EB3-Tomato, we further visualized MT polymerization into dendritic spines in live-cell imaging experiments (Figure 2J, K and Movie S1). Consistent with our former data, quantification confirmed a significant reduction of MT entries into spines with heads, following inhibition of katanin function by p60DEID (Figure 2L-N and Figure S3I), whereas spine density in general remained unaltered in p60DEID-expressing neurons (Figure S3J). Since the polymerization of MTs into dendritic spines occurs in a neuronal activity-dependent manner (22, 23), we combined EB3 immunostaining with the chemical induction of long-term potentiation (cLTP) (35). These experiments further confirmed that inhibition of katanin function (p60DEID) causes a significant reduction in the percentage of EB3-positive spines, as compared to control conditions (Figure 2O-S).

Taken together, we conclude that functional katanin participates in the polymerization of MTs into dendritic spines.

### Katanin regulates neuronal function and plasticity at potentiated synapses

Dendritic spine synapses are fundamental signaling units that mediate communication between individual neurons within neuronal circuits, and process information through long-term modification of their strength and structure. We therefore asked whether interference with microtubule severing (27), might alter synaptic transmission. An analysis of AMPAR-mediated miniature excitatory postsynaptic currents (mEPSCs) under basal conditions revealed comparable mEPSC amplitudes in the presence or absence (p60DEID) of functional katanin (Figure 3A, B). Likewise, the inter event intervals, were equal under both conditions (Figure 3A, C), indicating that basal synaptic transmission is independent of functional katanin.

**Figure 3.**
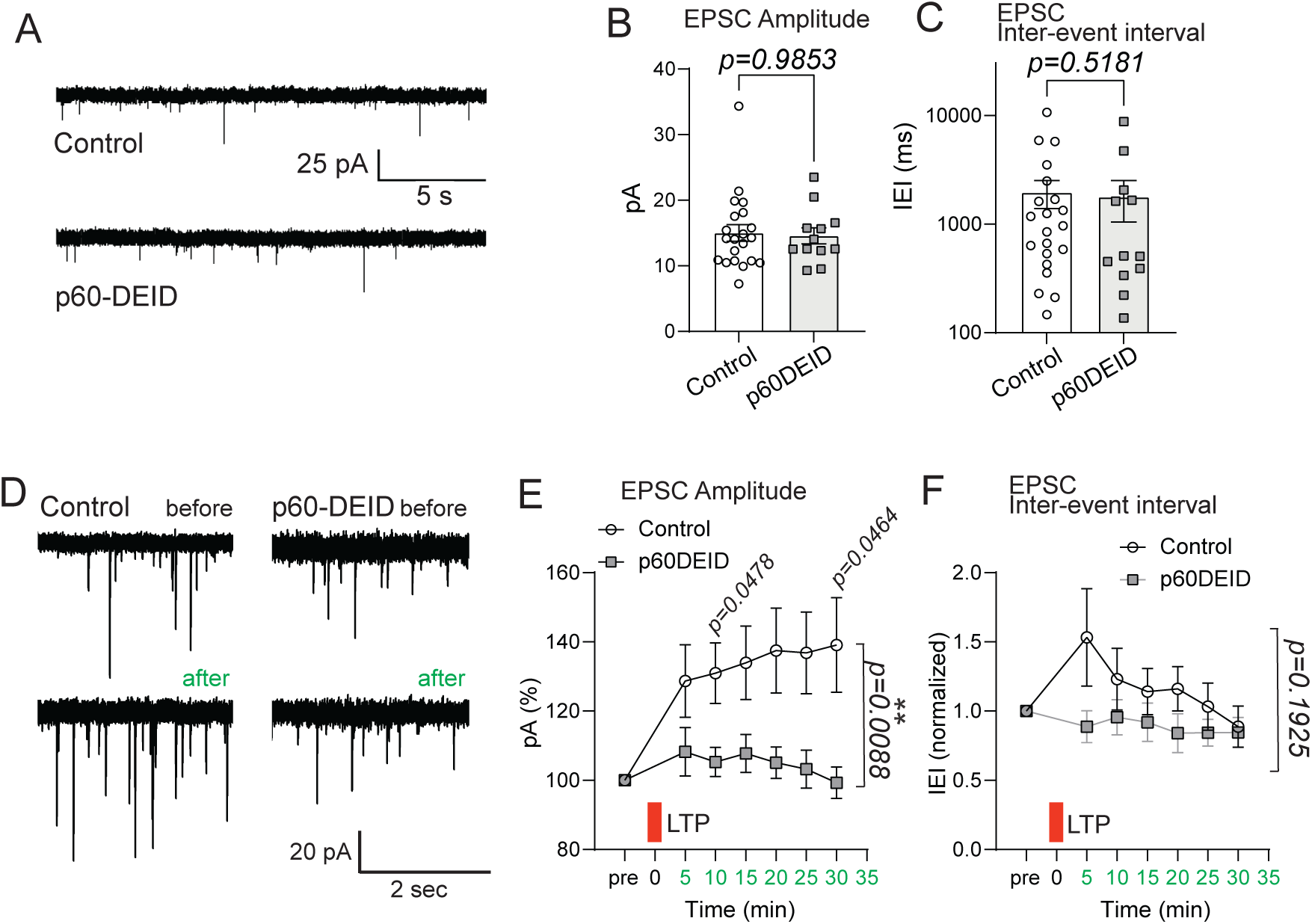
Katanin depletion or functional inhibition alters synaptic transmission. (A) Katanin p60DEID overexpression in primary hippocampal neurons does not alter mEPSCs under basal conditions, control n = 21 neurons, p60DEID n = 12 neurons. (B) Mean amplitudes (control: 15.03±1.26 pA; p60-DEID: 14.57±1.21 pA, Mann Whitney test p = 0.9853. (C) Medians of inter event intervals. Control: 1952±565.2 ms; p60-DEID: 1780±736.3 ms, Mann-Whitney-U-test p = 0.5181. (D) Katanin p60DEID overexpression in primary hippocampal neurons suppresses synaptic potentiation (cLTP). (E) EPSCs amplitudes. Stimulation increases the amplitudes of control transfected cells. Control, directly after cLTP induction: 128%; control, 30 min after cLTP induction: 139%; p60DEID, directly after cLTP induction: 108%; p60DEID, 30 min after cLTP induction: 99%. 2-way ANOVA: time*condition interaction p=0.0088. Individual data points t-tests: 10 minutes: p = 0.0478; 30 minutes: p = 0.0464. 5-minutes bins. (F) Inter-event intervals. 2-way ANOVA: time*condition no interaction p = 0.1925. Bins: 5 min. Green bar: cLTP stimulus.

Since MT invasion into dendritic spines is neuronal activity-dependent and increases, following NMDAR activation and calcium influx (22, 23), we further asked whether an induction of synaptic potentiation might depend on katanin-mediated functions. Activation of NMDAR-mediated synaptic transmission through removal of Mg^2+^ and application of glycine (LTP) (36) significantly increased mEPSC amplitudes, but not frequency (inter-event intervals) (Figure 3D-F, white circles). Notably, functional inhibition of katanin’s ATPase activity (p60-DEID overexpression) completely prevented this increase of mEPSC amplitudes following glycine application (Figure 3D-F, grey squares). Thus, katanin-mediated microtubule severing is a prerequisite for synaptic potentiation in primary neurons.

### Katanin regulates glutamate-induced structural spine remodeling

The structural remodeling of dendritic spines is driven by their dynamic actin cytoskeleton (37, 38) however to what extent katanin-mediated functions and/or microtubule entry (9, 15) contributes to spine structure is incompletely understood. Glutamate can be sufficient to trigger dendritic spine growth from dendrite shafts a process that requires opening of NMDARs and activation of cAMP-dependent protein kinase (PKA) (39).

To further investigate whether neuronal activity-induced spine remodeling (37, 38) requires functional katanin we applied two-photon glutamate uncaging at single spines on hippocampal neurons that expressed td-Tomato as a volume marker. Following stimulation of caged glutamate at single spines, we observed robust spine growth over time in control neurons (Figure 4A, B, upper and 4C, white circles). We then combined this assay with overexpression of katanin p60DEID to interfere with its microtubule-severing function (27). Notably, in the presence of the dominant-negative katanin mutant, the glutamate uncaging-induced growth of spine protrusions was strongly inhibited (Figure 4A, B, lower and 4C, grey squares), suggesting that katanin-mediated functions contribute to the activity-dependent remodeling of spines.

**Figure 4.**
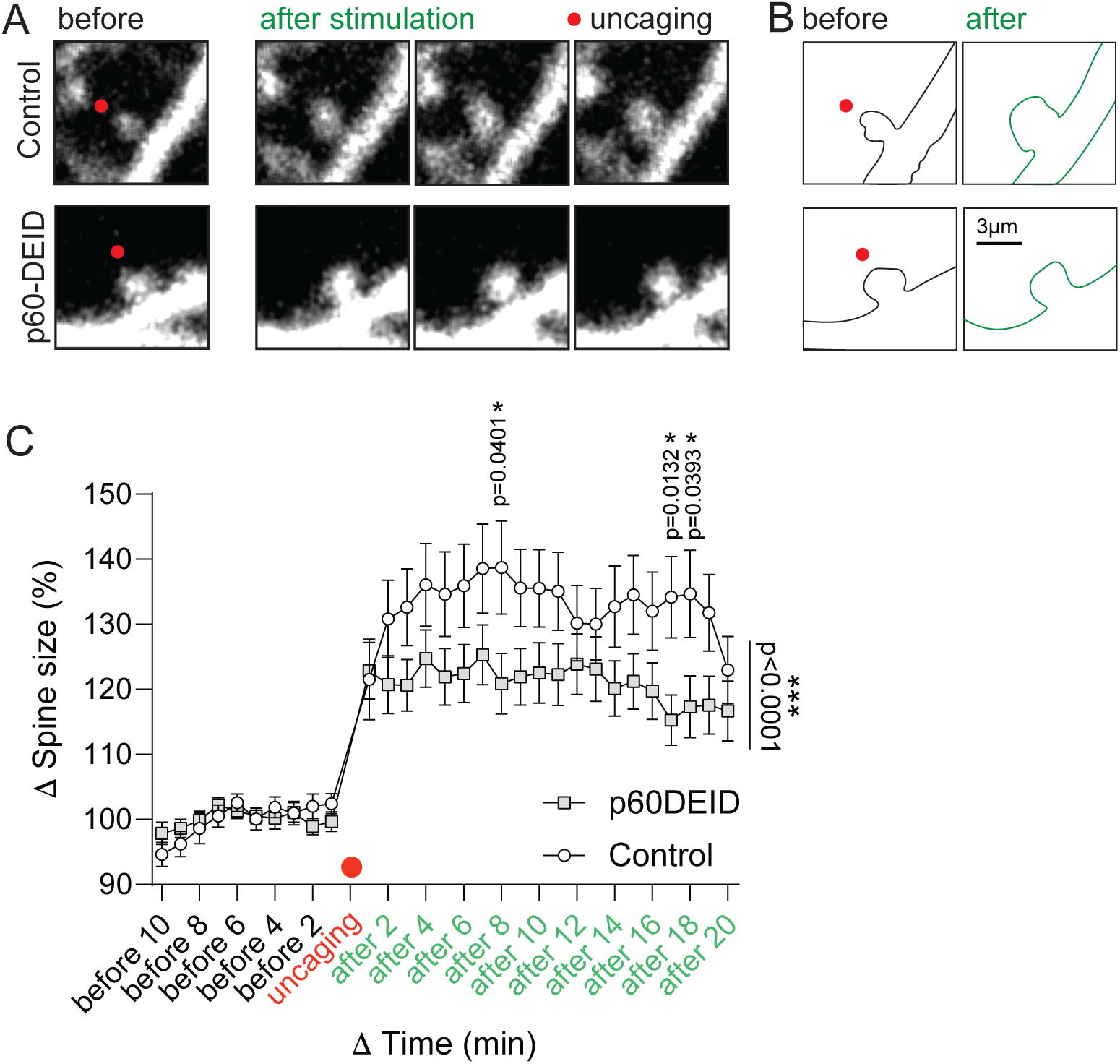
Inhibition of katanin function interferes with glutamate-induced structural spine remodeling. (A) Glutamate uncaging at single synapses using cultured hippocampal neurons. Left: before stimulation (black), right: after stimulation (green). Red dot: uncaging stimulus. (B) Schematic representation of spine size before (left, black) and after (right, green) glutamate stimulation (red dot). (C) Quantification of D shows a significant difference in spine size (Δ spine size) between control and p60DEID after stimulation. 2-way ANOVA test, significant interaction between time*condition, p<0.0001, multiple comparison with Fisher’s LSD test shows “after 8” p = 0.0401; “after 17” p = 0.0132; “after 18” p = 0.0393 (Control, n = 39; p60DEID n = 45), bins 2 min. Before uncaging 0-10 min (black axis labels), after uncaging 0-20 min (green axis labels).

Our data highlight katanin as a relevant regulator of structural synaptic plasticity, in particular under conditions of synaptic potentiation.

## Discussion

Here, we show that katanin is a synaptic protein complex mediating critical roles in regulating synapse structure and function under activity-dependent conditions.

Dendritic spines contain a prominent actin cytoskeleton that undergoes growth and shrinkage depending on glutamatergic transmission and its downstream signals (37, 38). In contrast to actin filaments being a permanent constituent, MTs transiently polymerize into and out of dendritic spines. This phenomenon is activity-dependent and requires NMDAR activation and calcium influx (19, 20, 40). For instance, chemical protocols to induce LTP and LTD increase or decrease these events, respectively (22–24). Furthermore, actin remodeling at the base of spines enables MT entry, thereby linking synaptic activity and actin remodeling with MT invasion into spines (9, 23, 41). A key study further identified that transient MT invasions mediate the transport of KIF1A and synaptotagmin IV into spines (21), suggesting a potential role in synaptic transport. However, whether MTs and/or their interacting proteins associate with the postsynaptic density is incompletely understood. Interestingly, the MT +TIP protein EB3 binds to the glutamate receptor scaffold protein PSD-95 (42). Our co-IP experiments (Figure 1J, K), which report association of a katanin subunit with PSD-95, are in line with this view. We therefore asked whether the MT severing protein katanin might regulate synaptic structure and/or function.

Within the hexameric katanin severing complex, the catalytic p60 subunit can be replaced by two alternative p60-like subunits (2, 43). Since genetic triple knockouts of p60 and its homologues are unavailable, we alternatively applied functional inhibition of katanin’s ATPase activity with a dominant-negative approach that has been shown to inhibit MT severing (27).

Using cultured primary neurons, we first analyzed microtubule dynamics in dendrites. At stage DIV4, when dendrite outgrowth occurs, but neurons do not yet contain dendritic spine protrusions (33), katanin inhibition altered MT growth velocities (Figure S3), suggesting that katanin-mediated functions might contribute to dendritogenesis at early developmental stages. This result is in line with former experiments injecting a function-blocking katanin-antibody into neurons that led to reduced neurite outgrowth (44). However, at stages DIV12-17 when neurons have developed dendritic spines (Figure 1, 2, S1, S2) and are characterized by functional EPSCs (Figure 3), we no longer observed changes in either MT growth velocity or EB3 comet density, suggesting that under basal conditions functional katanin is not critical in dendrites of mature neurons.

However, at these mature neuronal stages our data show that katanin participates in the activity-dependent invasion of MTs into individual dendritic spines (Figure 2). In general, MT severing is important to keep MTs in a dynamic state, for instance, to trigger neurite branching (26, 45–47). MTs within neurites are relatively long, but undergo local severing at branch points. As a result, shorter MTs are thought to develop new plus ends to polymerize into the newly forming branch. It has been unclear whether a similar local severing mechanism also occurs at the base of dendritic spines and induces polymerization of MTs into spines. Since dendritic spines are oriented perpendicular to the dendrite, local severing might be critical as it is at dendritic branch points (25). That katanin subunits are located at dendrites and spines (Figure 1) and that EB3-labeled microtubule +TIPs in spines are decreased upon inhibition of katanin function (Figure 2), support this view. The bending capacity of MTs might help them to enter the spine neck, although this is limited to about 1.7 rad/µm (48). In this respect, the removal of tubulin subunits by katanin (47), could increase MT flexibility or create new locations for branching. Finally, direct MT branching through a SSNA1-mediated mechanism (49) might regulate dendritic MTs.

Although MT polymerization into spines (19, 20) is activity-dependent (22, 23) and promotes kinesin-mediated cargo transport towards the postsynaptic specialization (21), it was unclear whether MT invasion has a functional and/or structural role at glutamatergic spine synapses. We found that katanin inhibition, which interferes with MT invasion into individual spines, significantly reduces LTP of EPSC amplitude (Figure 3) and reduces glutamate-uncaging induced structural spine plasticity (Figure 4). Our data suggest that katanin-mediated MT severing might act upstream of MT polymerization into dendritic spines. They further support the view that, in addition to actin, MTs are important for synaptic function.

## Materials and Methods

For detailed materials and methods, see Supplementary information.

### Animals

Maintained in the animal facility of UKE in accordance to European Communities Council Directive (2010/63/EU). Tissue harvesting in accordance to the Hamburg ethics committee (Behörde für Justiz und Verbraucherschutz, Fachbereich Lebensmittelsicherheit und Veterinärwesen).

### Cell fractionation and co-immunoprecipitation

For S1-S2 and P2 fractions as well as for synaptosomal preparations, hippocampi of adult mice were used. Co-immunoprecipitation was achieved by harvesting forebrains of adult mice. Protein G dynabeads (1004D, Invitrogen) were incubated with p80 katanin or PSD-95 antibodies or rabbit IgG as a control.

### Primary cell cultures

Hippocampi were harvested at E.16. Neurons were plated at different densities. Transfection was achieved either with calcium phosphate or Lipofectamine 2000 protocols.

### Immunocytochemistry

Neurons were used for experiments at DIV 12-17. Depending on the experiment, neurons were fixed with 4% Formaldehyde/PBS or (for EB3 analysis in spines) ice-cold methanol followed by post-fixation with 4% Formaldehyde/PBS. Whenever applicable, neurons were transfected two days before fixation. For endogenous EB3 labeling upon stimulation, glycine/no Mg^2+^ was administered 2 days after transfection.

### Time-lapse imaging

Hippocampal neurons were co-transfected at DIV 12. Two days later, neurons were observed at a spinning disk microscope. 300 frames with an interval of 2 seconds were acquired sequentially for GFP and RFP channels. Fiji (NIH, Version 2.0.0-rc-23/1.49m) was used for visualization and quantification. Processing of images was identical throughout experiments.

### Two-photon glutamate uncaging

DIV 12-17 hippocampal neurons were transfected and, after 48 hours, were observed under an Olympus FV1000 microscope equipped with Fluoview software. Ti: sapphire laser (Mai Tai, Spectra Physics) tuned to 880 or 950 nm was used for excitation. Images were collected with an Olympus XLPlanN 25X MP, 1.05 NA water dipping objective and a 495-540nm emission filter. Glutamate was uncaged with laser set to 720nm with 60 pulses, 1ms at 1 Hz.

## Acknowledgements

We thank F. J. McNally for the p60DEID construct and E. Szpotowicz and C. Raithore for technical assistance. Supported by DFG grants KN556/11-2 (FOR 2419) and KN556/12-1 to MK, GE2762/3-2 (FOR 2419) to CGE, by the City of Hamburg grants LFF-FV74 to MK and MG and by LFF-FV76 to MK.

## Supplementary Information

**Figure S1 (related to Figure 1).**
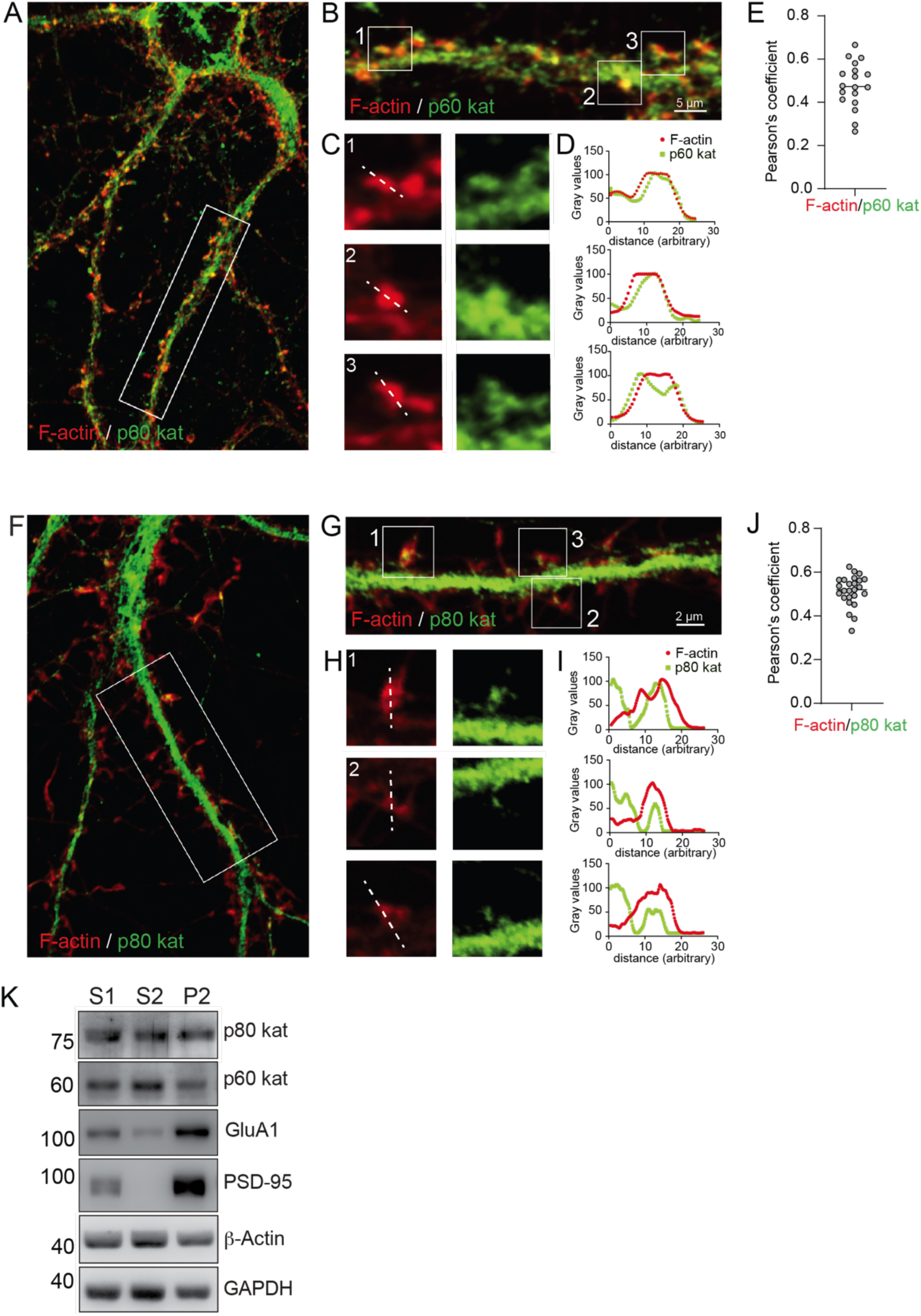
Katanin is located at F-actin-enriched spine compartments. (A) Immunostaining of endogenous katanin p60 (green), Rhodamine-phalloidin labeling of F-actin (red), DIV13-17 neurons, N = 3 experiments. (B) Boxed region in A. (C) Magnifications of spines from boxed regions in B. (D) Fluorescence intensity profiles of line scans in C. (E) Pearson’s correlation coefficient indicating colocalization between p60 katanin and F-actin. Mean±S.E.M = 0.4797±0.02536, n = 18 ROIs. (F) Immunostaining of endogenous katanin p80 (green), Rhodamine-phalloidin labeling of F-actin (red), DIV13 neurons, N = 3 experiments. (G) Boxed region in F. (H) Magnifications of spines from boxed regions in G. (I) Fluorescence intensity profiles of line scans in H. (J) Pearson’s correlation coefficient indicating colocalization between p80 katanin and F-actin. Mean±S.E.M = 0.5149±0.01379, n = 25 ROIs. (K) Western blot detection following differential centrifugation. GAPDH: loading control. S1: supernatant 1,000 x g fraction, S2: supernatant 10,000 x g fraction, P2: Plasma membrane-enriched pellet 10,000 x g, (n = 3).

**Figure S2 (related to Figure 1).**
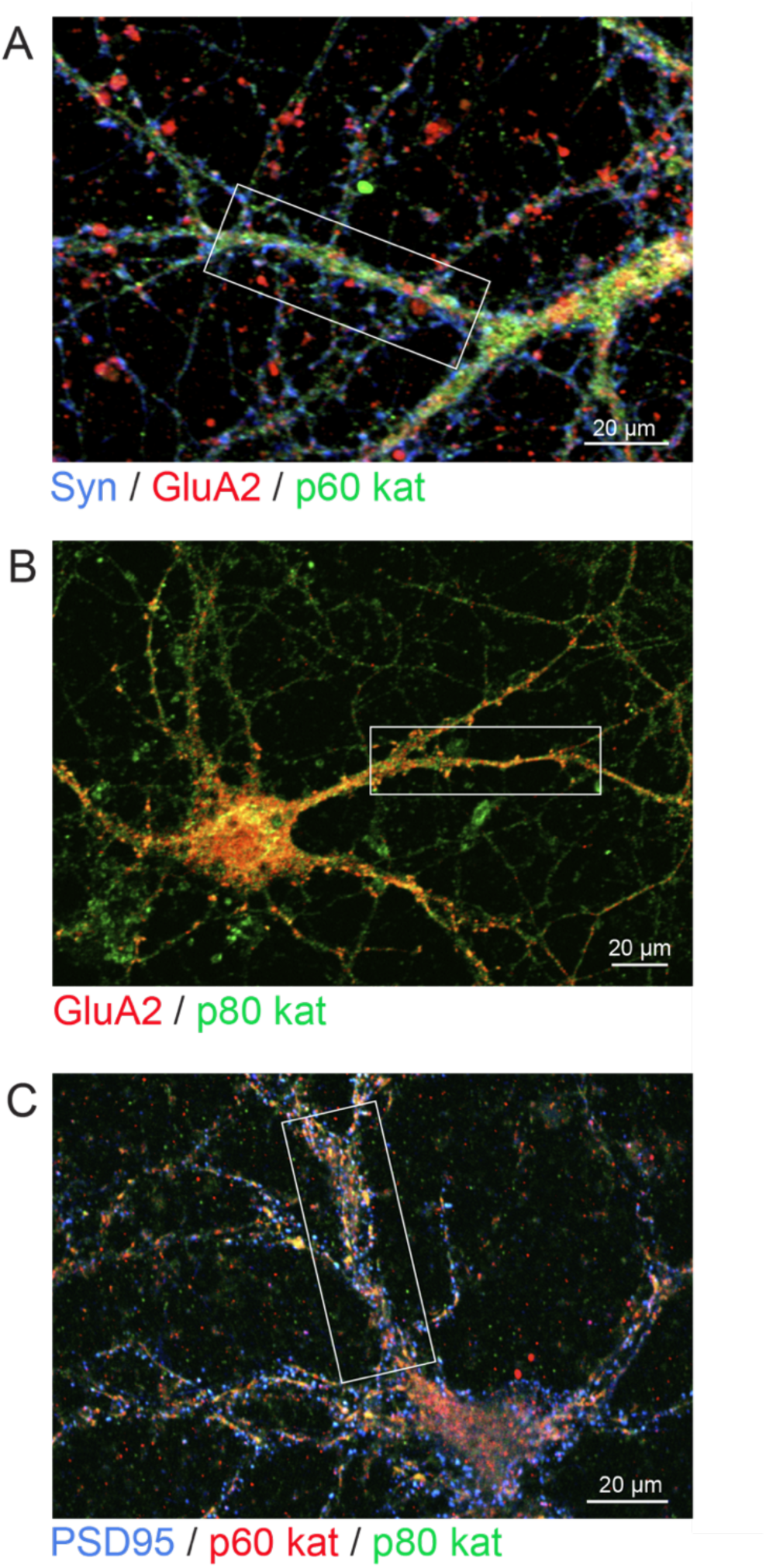
Colocalization of katanin subunits with pre- and postsynaptic marker proteins. Immunostaining of endogenous proteins using primary hippocampal neurons in culture. (A) Synaptophysin (blue), AMPAR GluA2 (red), p60 katanin (green), related to Figures 1 A-D. (B) AMPAR GluA2 (red), p80 katanin (green), related to Figures 1 E-H. (C) PSD95 (blue), p60 katanin (red), p80 katanin (green), related to Figures1 L-M.

**Figure S3 (related to Figure 2).**
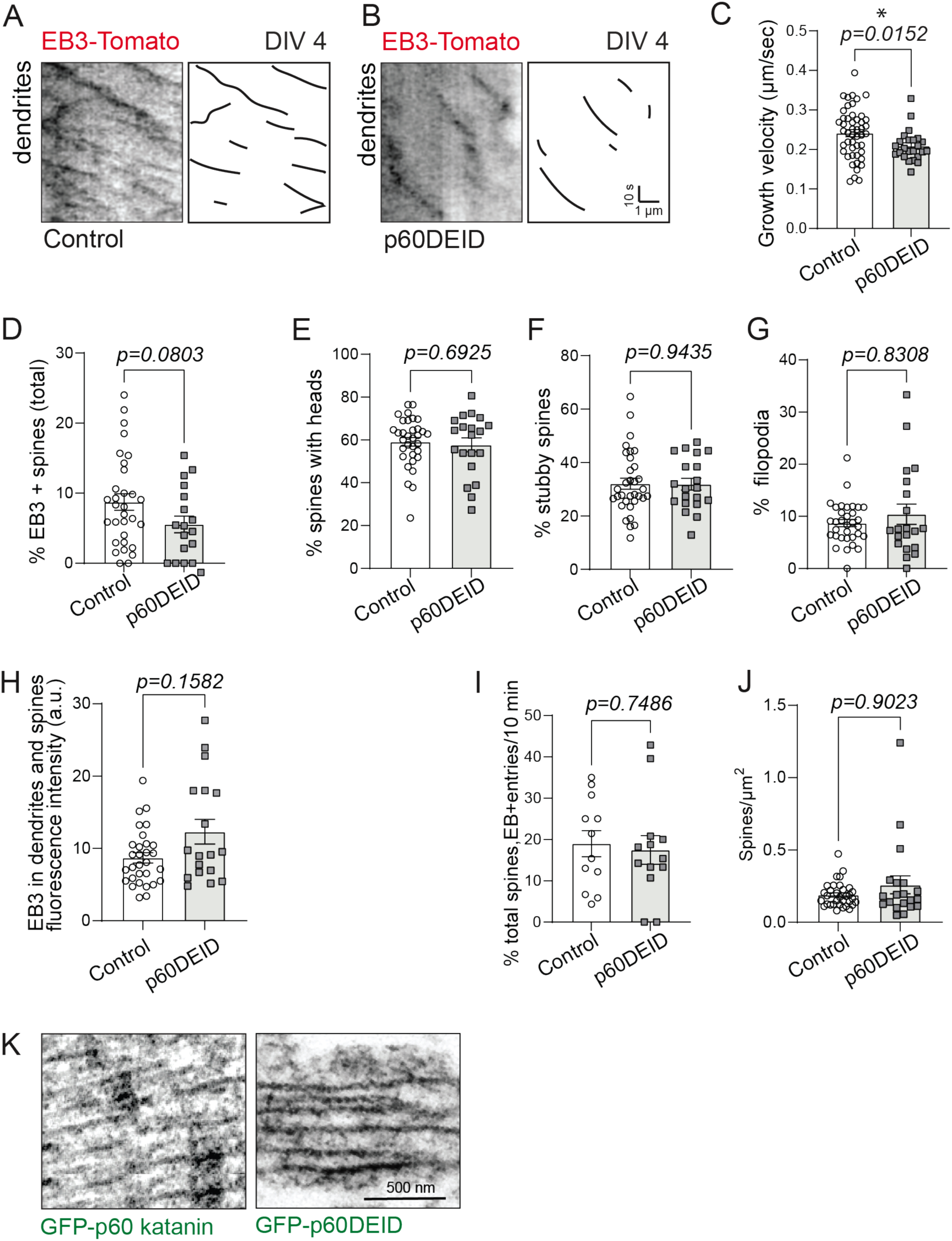
Functional inhibition of katanin reduces MT polymerization into dendritic spines. (A, B) EB3-Tomato time-lapse imaging in dendrites of DIV4 hippocampal neurons Right: cartoons of EB3 tracks for control in A and for p60DEID in B. (C) Quantification of EB3 growth velocity (µm/s). Control: 0.24±0.01, p60DEID: 0.21±0.01. Mann-Whitney-U-test p = 0.0152. Control n = 48 comets; p60DEID n = 24 comets. (D) Quantification of %EB3- positive total spines. Control: 8.74±1.16%, p60DEID: 5.55±1.19%. Unpaired two-tailed t-test p = 0.0803. (Control: n = 33 cells, 1621 spines; p60DEID: n = 20 cells, 648 spines). (E) Quantification of % spines with heads. Control: 59.16±2.01%, p60DEID: 57.72±3.23%, unpaired two-tailed t-test p = 0.6925 (Control: n = 33 cells, p60DEID: n = 20 cells). (F) Quantification of % stubby spines. Control: 32.10±2.08%, p60DEID: 31.88±2.19%, unpaired two-tailed t-test p = 0.9435 (control: n = 33 cells, p60DEID: n = 20 cells). (G) Quantification of % filopodia. Control: 8.70±0.70%, p60DEID: 10.40±1.95%, Mann-Whitney-U-test p = 0.8308 (Control: n = 33 cells, p60DEID: n = 20 cells). (H) EB3 fluorescence intensity in dendrites of control and p60DEID transfected neurons N = 3 experiments; Control: 8.70+-0.70; p60DEID: 12.31+-1.70. Mann-Whitney-U-test p = 0.1582. (Control = 33 cells, p60DEID = 20 cells). (I) Percentage of total spines invaded by EB3-Tomato within 10 min: control: 19.00±3.15%, p60DEID: 17.48±3.45%. Unpaired two-tailed t-test p = 0.7486. (Control: n = 12 cells; p60DEID: n = 13 cells). (J) Quantification of spine density (number of spines/µm^2^). Control: 0.19±0.01%, p60DEID: 0.26±0.06%. Mann-Whitney-U-test p<0.9023. (Control: n = 33 cells, p60DEID n = 20 cells). (K) Anti-GFP immunoelectron microscopy with diaminobenzidine (DAB) showing MTs from hippocampal neurons transfected with either EGFP-p60 katanin or EGFP-p60DEID.

**Movie S1.**
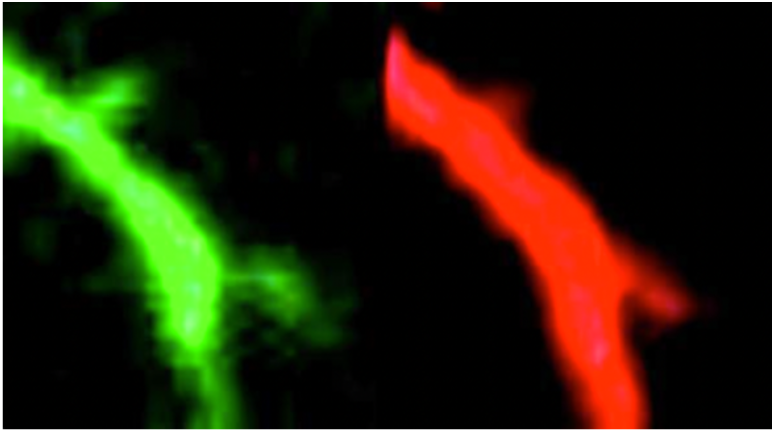
Still image of supplemental movie to Figure 2J. An EB3-positive signal representing microtubule polymerization is identified in the lower spine protrusion over time. The movie shows the validity of the method. Acquisition rate: 1 frame/s.

## Detailed Materials and Methods

### Animals

Animals were maintained in the animal facility of the University Medical Center Hamburg-Eppendorf under controlled environmental conditions in accordance with the European Communities Council Directive (2010/63/EU). Harvesting of tissue and cells was performed according to the Hamburg authorities (Behörde für Justiz und Verbraucherschutz, Fachbereich Lebensmittelsicherheit und Veterinärwesen) and the animal care committee at the University Medical Center Hamburg-Eppendorf.

### Cell Culture and Transfection

Hippocampal neurons were cultured as following: E.16 pregnant female mice were sacrificed and their embryos harvested from the uterine horns. Embryos were decapitated and their hippocampi dissected under the microscope. Hippocampi were subjected to 0.05% trypsin for 5 min at 37°C, followed by mechanical dissociation using a glass Pasteur pipette. Neurons were plated at different densities around 70,000 cells per coverslip. Transfection was performed using the calcium phosphate method (1, 2) or using Lipofectamine 2000.

### Immunocytochemistry

Primary cultured hippocampal neurons between DIV12 and DIV17 were fixed for 4-5 min in 4% Formaldehyde/PBS and washed 3 times in PBS for 5 min each. Cells were permeabilized for 5 min using 0.2% Triton X-100/PBS, followed by three washes of 5 min each with PBS. Blocking was performed for 1 hour at room temperature with 1% BSA/PBS. Primary antibodies were incubated in 1% BSA/PBS for 1 hour at room temperature. After three washes with PBS as before, secondary antibodies were incubated for 1 hour in 1%BSA/PBS with or without Rhodamine/Phalloidin 1:500. Mounting was performed after three washes. *EB3 staining*: hippocampal neurons were transfected between DIV12 and DIV17 and the experiment was conducted 48 hours later.

Cultures were fixed with ice-cold methanol for around 4 min, washed once with PBS at room temperature for 20 min and post-fixed in 4% Formaldehyde /PBS for additional 4 min. Next, two washes with PBS were performed for a total time of 30 min, followed by blocking for 45 min at room temperature with 1% BSA/PBS. Primary antibodies were incubated overnight in a wet chamber diluted in Phem Buffer (60mM Pipes, 25 mM HEPES, 10 mM EGTA, 2 mM MgCl_2_, pH=6.9). After, three washes with PBS, secondary antibodies were diluted in Phem Buffer and incubated at room temperature for 2 hours. Finally, three washes with PBS were performed followed by mounting with Aqua Poly Mount 1. Imaging was achieved using an Olympus FV1,000 confocal microscope. Images were quantified using Fiji (NIH, Version 2.0.0-rc-23/1.49m). Spines that contained at least 1 fluorescent pixel were considered positive for EB3. When EB3 puncta were found partially inside spines, if the majority of puncta or the pixels of highest value were inside, the spine was considered as positive. Based on their general morphology, spines were classified into spines with heads, stubby spines and filopodia. *Glycine stimulation for EB3 staining*: hippocampal cultures were transfected at DIV12 as above. Stimulation was performed for 3-5 min in Mg2+-free ACSF containing 50mM Bicuculline, 1µM TTX and 200 mM glycine (3). Subsequently, cells were recovered in Mg2+-free ACSF for either 30-60 min. Immunostaining was performed as described above. Spine classification was performed based on their general morphology into spines with heads, stubby spines and filopodia.

### Immunoelectron microscopy

Pre-embedding immunolabelling was performed as following: dissociated neurons were fixed with PB buffer containing 4% paraformaldehyde and 0.1% glutaraldehyde. 2.3 M sucrose was used for cryoprotection. Penetration of immunoreagents inside the cells was obtained by two freeze-thaw cycles in liquid nitrogen. After washing in PBS, 10% horse serum (PS) containing 0.2% bovine serum albumin (BSA) was used for blocking for 15 min. Primary antibodies were incubated overnight in PBS with 1% PS and 0.2 % BSA (Carrier). Cells were washed with PBS and treated with biotinylated secondary antibody diluted in Carrier solution for 90 min. Subsequently, cultures were washed and incubated with ABC (Vector Labs) 1:100 in PBS for 90 min. This was followed by washing and incubation in diaminobenzidine (DAB)-H202 solution (Sigma St. Louis, USA) for 10 min. Thereafter, cultures were washed three times in 0.1 M sodium cacodylate buffer (pH 7.2–7.4) followed by osmication with 1% osmium tetroxide in cacodylate buffer. Dehydration using ascending ethyl alcohol concentration steps was followed by two rinses in propylene oxide. The embedding medium was infiltrated by first immersing the sections in a 1:1 solution of propylene oxide and Epon followed by neat Epon. Finally, samples were hardened at 60 °C. 0.5 µm sections (semithin) from the hippocampal region were mounted on glass slides and stained for 1 minute with 1% Toluidine blue. 60 nm sections (ultrathin) were observed with a JEM-2100Plus Transmission Electron Microscope at 200kV (Jeol, Germany). Images were acquired with the XAROSA CMOS camera (Emsis, Germany).

### Cell fractionation and Co-immunoprecipitation

#### S1-S2 and P2 membranous fraction

Hippocampi of 3 *wildtype* mice at 14 weeks old were harvested and treated separately on ice. Briefly, hippocampi from one mouse were suspended in sucrose 1 buffer (320mM sucrose, 1mM NaHCO_3_, 1mM MgCl_2_, 500μM CaCl_2_, EDTA-free protease inhibitors). Hippocampi were homogenized with 12 potter strokes at 900 rpm and centrifugated at 1,400 x g for 10 min at 4°C. An aliquot was saved for further analysis (S1). The rest was centrifugated at 13,800 x g for 10 min at 4°C. The supernatant (S2 fraction) was saved and the pellet (P2-membranous) was resuspended in sucrose 2 buffer (320mM sucrose and 1mM NaHCO_3_). All fractions were quantified using BCA Protein Assay kit (23227, Thermo Scientific) and normalized to equal concentrations using sucrose buffers 1 and 2, when appropriate. *Post-synaptic density isolation*. Crude membranous fractions from hippocampi (P2) were prepared as above. Hippocampi were isolated and homogenized at 900 rpm with ten strokes. Homogenates were centrifugated at 4°C for 10 min at 1,000 x g. Supernatants were conserved, pellets resuspended in sucrose buffer 1 and centrifugated at 700 x g for 10 min at 4°C (JA25.5 rotor). Combined supernatants were centrifugated at 13,800 x g for 10 min at 4°C to obtain P2 fractions, which were resuspended in sucrose buffer 2. In parallel, a sucrose gradient (0.85M, 1M and 1.2M plus 1mM NaHCO3) was prepared in Ultra-clear 14x95mm, 14 ml tubes. The P2 fraction was placed on top of the gradient and centrifugated for 90 min at 82,500 x g and 4°C with a SW40-Ti rotor. Synaptosomal layers appeared between 1M and 1.2M. They were diluted 4-fold in sucrose buffer 2 and centrifugated at 28,000 x g for 20 min at 4°C using a JA 25.50 rotor. Pellets (synaptosomes) were resuspended in 100 microliters of sucrose 2 buffer. For PSD fractions, synaptosomes were diluted with 2x Triton X-100 buffer and shaked for 15 min. The Triton-X-100-soluble fraction was separated from the Triton-X-100-insoluble fraction via centrifugation at 70,000 x g and 4°C for 1 hour. Pellets were dried for 15 min and then resuspended in Tris-HCl pH=8.0.

#### Immunoprecipitation

Forebrains of 3 *wildtype* mice were harvested and treated independently. Briefly, S1-S2 and P2 fractions were prepared as above. Subsequently, protein G dynabeads (1004D, Invitrogen) were washed twice with IP wash buffer (20mM HEPES, 100mM potassium acetate, 40mM KCl, 5mM EGTA, 5mM MgCl_2_, pH=7.2) containing 10% Triton X-100. Beads were incubated with 6μg of either rabbit IgG (control), p80 katanin or PSD-95 primary antibodies for 1.5 hours (Ab-beads). Ab-beads were washed twice with IP wash buffer containing Triton-X-100 and incubated with forebrain P2 fractions overnight at 4°C (Ab-beads-lysates). The following day, Ab-beads-lysates were washed six times in IP wash buffer. After washing step number 3, an intermediate incubation in IP wash buffer was performed on a rotating wheel. Finally, beads were resuspended in 200 μl SDS-Loading buffer and boiled. The P2 fraction in SDS-loading buffer was considered as input. A western blot was carried out in the presence of either 2% BSA or IgG-free BSA.

Samples were processed by SDS PAGE and subsequently blotted onto a PVDF membrane. Transfer was achieved in the presence of 20% methanol transfer buffer (192mM glycine, 25mM Tris). Blocking was performed in 5% milk TBS-T. Antibodies were incubated either in 5% BSA, TBS-T or 5% milk, TBS-T.

### Time-lapse Video Microscopy, Image Processing

Neurons were imaged with a spinning disk microscope system (Visitron Systems GmbH), consisting of a Nikon Ti-E inverted microscope and a Yokogawa spinning disk. A chamber for temperature (37°C) and CO_2_ (5% CO_2_) control was used. *EB3 in dendrites of young neurons.* Hippocampal cultures at DIV3 were co-transfected with EB3 tomato and either p60DEID or GFP. Neurons were imaged at DIV4. A single acquisition was obtained for the GFP channel, while for the EB3-Tomato channel, time lapse images of 1 frame per second were taken. Quantification was performed with Fiji (NIH, Version 2.0.0-rc-23/1.49m) using manual tracking plugin. No difference was made between anterograde and retrograde movement and the focus was in neuronal dendrites. Absolute speed (no stops), was considered for analysis. *EB3 in dendritic spines.* Hippocampal neurons at DIV12 were co-transfected with EB3-Tomato and either EGFP (control) or EGFP-p60-DEID. Briefly, eight microliters of lipofectamine (Lipofectamine 2,000, Invitrogen) were incubated in 50 microliters Opti-MEM. In parallel, a total of 4-8 micrograms of DNA were incubated in 50 microliters Opti-MEM. After 5-10 min, both reactions were mixed and administered to the transfected coverslip, after removal of around 75% of the conditioned media. 2 hours after incubation, transfection was aspirated and replaced by conditioned media without washing. At DIV14, transfected neurons were imaged in Hepes media. Imaging was performed sequentially for green and red channels with an interval of 2 seconds between frames for a total of 300 frames. Binning of 2 was used for image acquisition. Quantification was performed using Fiji (NIH, Version 2.0.0-rc-23/1.49m). Images were processed identically throughout experiments. The GFP channel was used to check for the expression of either EGFP-p60-DEID or control. The RFP channel was used for quantification. Brightness and contrast were adjusted in order to observe neuronal morphology due to the non-comet EB3 diffusion (4). Individual ROIs were created and saved to mark every possible spine. Spines were classified visually in spines with heads, stubby (local deformation of the dendrite) and filopodia (very motile, long filamentous). Back in the raw image, after adjusting brightness and contrast in the same way for all the images, ROIs of the identified spines were pasted and each spine was analyzed individually. The presence or absence of an EB3 comet was assessed visually. EB3 invasion into spines was considered positive only if the comet fully invaded the spine and ended its trajectory in the protrusion. The length of the events and the frame in which they appeared was also considered.

#### EB3 imaging in dendrites of mature neurons

DIV14 neurons imaged above were analyzed using Fiji (NIH, Version 2.0.0-rc-23/1.49m) and the TrackMate Plugin. Acqusition: 1 frame/2s. All tracks were manually checked after semi-automatic tracking.

### Patch-clamp recordings of EPSCs

Whole-cell patch clamp measurements (Hamill et al., 1981) were performed on DIV12-14 hippocampal neurons. Neuron cultures were prepared from embryonic (E16) mice. Recordings were done by using borosilicate pipettes with resistances of 3.0 - 4.5 MΩ after filling with intracellular solution (120 mM K-gluconate, 8 mM NaCl, 2 mM MgCl_2_, 0.5 mM CaCl_2_, 5 mM EGTA, 10 mM HEPES, 14 mM phosphocreatine, 2 mM magnesium-ATP, 0.3 mM sodium-GTP, and pH adjusted to 7.3 with KOH). Patchmaster software (HEKA, Lambrecht, Germany) in combination with an EPC-9 patch-clamp amplifier (HEKA) was used for data acquisition and pulse application.

Recordings were low-pass filtered at 2.9 kHz and analyzed with Fitmaster (HEKA, Lambrecht, Germany), Igor Pro 6.03 (Wavemetrics), Mini Analysis (Synaptosoft, Decatur, GA), and Excel (Microsoft). Neurons with an access resistance <20 MΩ were evaluated. Miniature excitatory postsynaptic currents (mEPSCs) were recorded from neurons transfected either with GFP (control neurons) or with a dominant-negative mutant of katanin (P60-DEID, test neurons) and GFP. Current clamp experiments were done at room temperature (21-23°C) in Ringeŕs solution (143 mM NaCl, 5 mM 1 KCl, 0.8 mM MgCl_2_, 1 mM CaCl_2_, 10 mM HEPES, 5 mM glucose, and pH adjusted to 7.3 with NaOH). mEPSCs were measured in the presence of TTX (0.25 μM), bicuculline (10 μM), and AP5 (20 µM), which were added to the Ringeŕs solution. All substances were purchased from Sigma Aldrich. Chemical LTP was induced with the following protocol: Neurons were preconditioned with the NMDA antagonist AP5 (50 µM) for 24 h before patch clamp experiments. After obtaining the whole cell configuration and recording of the resting potential and action potentials mEPSCs were recorded for 5 min at resting conditions. Thereafter neurons were stimulated with a series of 10 depolarizing pulses of 70 mV amplitude and 100 ms duration with intervals of 50 ms. During the electrophysiological stimulation the Ringeŕs solution was changed to a Mg^2+^-free solution containing 100 µM glycine and 3mM CaCl_2_ (Lu et al., 2001). Only experiments that were stable for 30 min after stimulation were evaluated. mEPSC amplitudes and inter-event-intervals (IEI) were determined by using Mini Analysis (Synaptosoft) and summarized in bins of 5 min.

### Two-photon glutamate uncaging

E16 embryos were used for the preparation of primary hippocampal neurons in culture. Cells were transfected with either EGFP-p60-DEID/mCherry or EGFP/mCherry (control) after DIV 12 using lipofectamine (Lipofectamine 2,000, Invitrogen) or a calcium phosphate protocol. After 24-48 hours of expression, 2-photon glutamate uncaging was performed on an Olympus FV1,000 microscope controlled by the FluoView software. Excitation was achieved by a pulsed Ti::sapphire laser (Mai Tai, Spectra Physics) tuned to 880 or 950 nm. 4 Z-stack images were collected with an Olympus XLPlanN 25X MP, 1.05 NA water dipping objective and a 495-540nm emission filter. Experiments were conducted in Mg^2+^ free Ringer solution (125 mM NaCl, 2.5 mM KCl, 2 mM CaCl_2_, 33 mM (D)-Glucose, 25 mM HEPES, pH 7.3) in the presence of 2.5 mM MNI-caged-L-glutamate (Tocris) and 0.001 mM TTX (Tocris). 2-4 dendritic spines were targeted for potentiation in each neuron. After imaging, a baseline of 5-7 time points, glutamate was uncaged by setting the laser to 720 nm with 60 pulses, 1 ms at 1 Hz. Following uncaging the spine volume was monitored for 15-20 min.

### Statistical analysis

Data exploration, statistical analysis and graphical representation were performed with GraphPad Prism 9.0.1. *EB3-tomato time-lapse imaging in young neurons.* EB3 comets were analyzed using the manual tracking tool of Fiji. Only comets observed in dendrites were considered for analysis. No discrimination was made between anterograde vs retrograde movement. The effective velocity was considered for statistical tests, therefore start points and stops (0.000 values) were removed. Data were explored using descriptive statistics and normal distribution. The Anderson-Darling test was used to assess for a normal distribution. The Two-tailed Mann-Whitney-U-test was used to assess significance. *Endogenous katanin and EB3 in dendritic spines.* The Anderson-Darling test was used to assess for a normal distribution. The Two-tailed Mann-Whitney-U-test was applied between conditions “no p60-no EB3” vs “yes p60-yes EB3”. *Endogenous EB3 in dendritic spines upon p60-DEID overexpression and EB3-tomato time-lapse in dendritic spines.* A parametric T-test was performed following assessment of a normal distribution. If data showed not normal distribution, a non-parametric Mann-Whitney-U-test was applied. *Patch-clamp recordings of EPSCs.* Outliers were excluded by the following link: https://www.graphpad.com/quickcalcs/Grubbs1.cfm, followed by

GraphPad exploration and analysis. For peak amplitudes, the percentage was calculated and normalized to the prior to stimulation time point. Significant interactions were observed between time and condition using 2-way ANOVA. For IEI, fold change was calculated with respect to the before stimulation time point (set to 1). *Two-photon glutamate uncaging.* Spine areas were marked by drawing freehand regions around the spine perimeter. To assess changes in spine size, the average of the area from individual spines during the baseline was set as 100%. Normalization was performed for each individual time point. Outliers were removed with SPSS. GraphPad was used to compute 2-way ANOVA. Significance was assessed by multiple comparisons with the uncorrected Fisher’s LSD test.

---

**Table S1.**
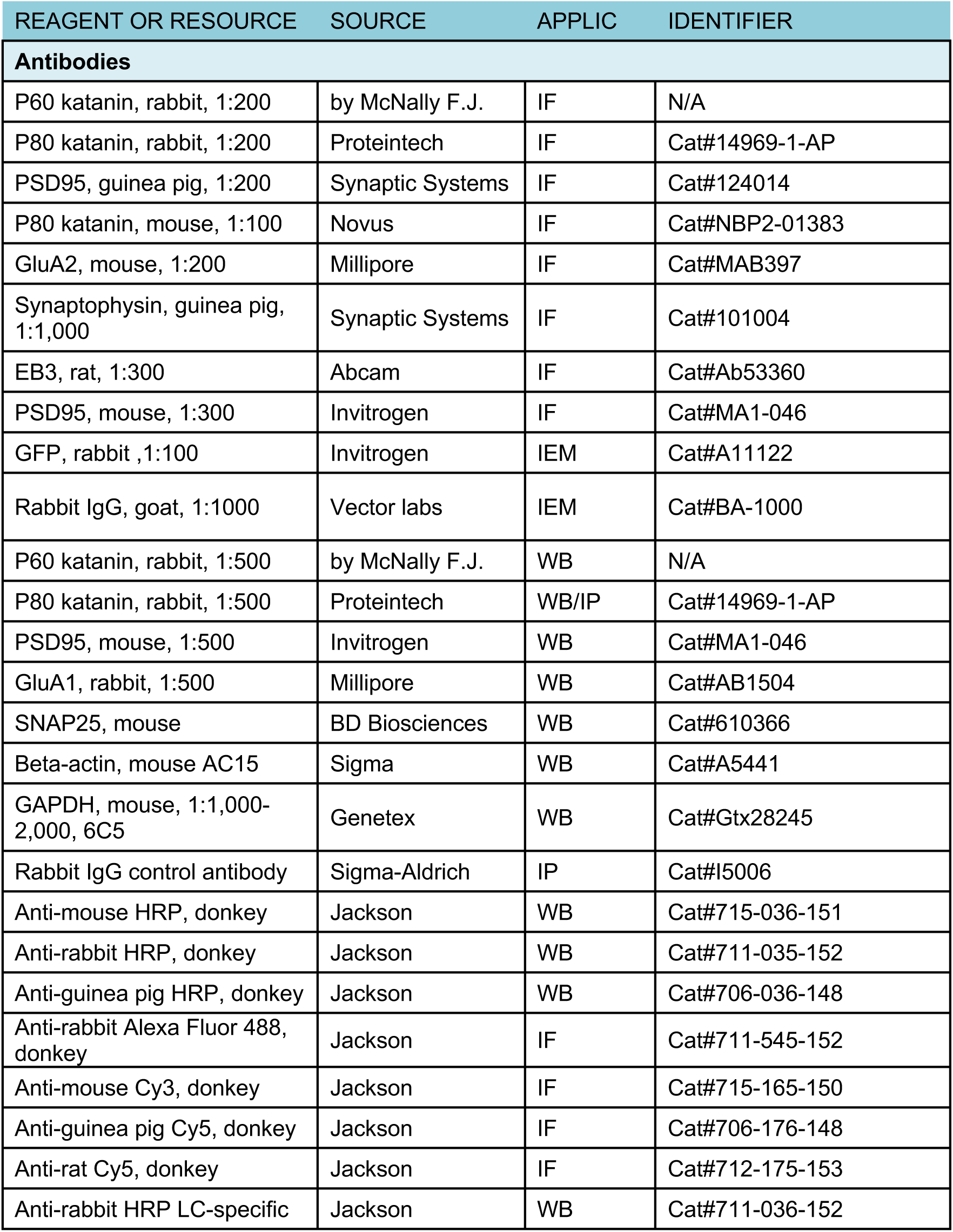
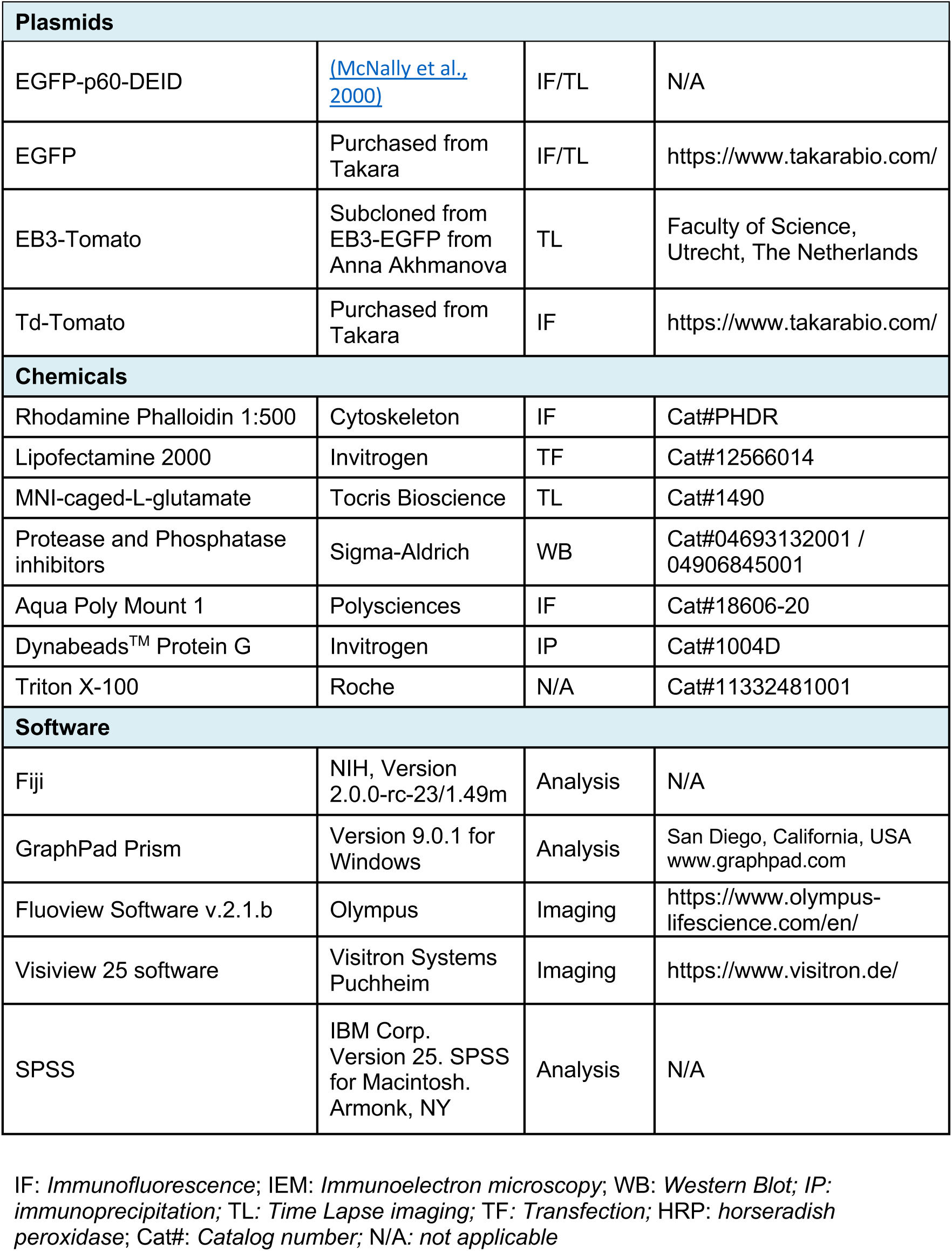
Antibodies, plasmids, chemicals and software.

## Notes

### Competing Interest Statement

The authors have declared no competing interest.

